# Design and proof-of-concept for targeted phage-based COVID-19 vaccination strategies with a streamlined cold-free supply chain

**DOI:** 10.1101/2021.03.15.435496

**Authors:** Daniela I. Staquicini, Fenny H. F. Tang, Christopher Markosian, Virginia J. Yao, Fernanda I. Staquicini, Esteban Dodero-Rojas, Vinícius G. Contessoto, Deodate Davis, Paul O’Brien, Nazia Habib, Tracey L. Smith, Natalie Bruiners, Richard L. Sidman, Maria L. Gennaro, Edmund C. Lattime, Steven K. Libutti, Paul C. Whitford, Stephen K. Burley, José N. Onuchic, Wadih Arap, Renata Pasqualini

## Abstract

Development of effective vaccines against Coronavirus Disease 2019 (COVID-19) is a global imperative. Rapid immunization of the world human population against a widespread, continually evolving, and highly pathogenic virus is an unprecedented challenge, and many different vaccine approaches are being pursued to meet this task. Engineered filamentous bacteriophage (phage) have unique potential in vaccine development due to their inherent immunogenicity, genetic plasticity, stability, cost-effectiveness for large-scale production, and proven safety profile in humans. Herein we report the design, development, and initial evaluation of targeted phage-based vaccination approaches against Severe Acute Respiratory Syndrome Coronavirus-2 (SARS-CoV-2) by using dual ligand peptide-targeted phage and adeno-associated virus/phage (AAVP) particles. Towards a unique phage- and AAVP-based dual-display candidate approach, we first performed structure-guided antigen design to select six solvent-exposed epitopes of the SARS-CoV-2 spike (S) protein for display on the recombinant major capsid coat protein pVIII. Targeted phage particles carrying one of these epitopes induced a strong and specific humoral response. In an initial experimental approach, when these targeted phage particles were further genetically engineered to simultaneously display a ligand peptide (CAKSMGDIVC) on the minor capsid protein pIII, which enables receptor-mediated transport of phage particles from the lung epithelium into the systemic circulation (termed “dual-display”), they enhanced a systemic and specific spike (S) protein-specific antibody response upon aerosolization into the lungs of mice. In a second line of investigation, we engineered targeted AAVP particles to deliver the entire S protein gene under the control of a constitutive cytomegalovirus (CMV) promoter, which induced tissue-specific transgene expression stimulating a systemic S protein-specific antibody response. As proof-of-concept preclinical experiments, we show that targeted phage- and AAVP-based particles serve as robust yet versatile enabling platforms for ligand-directed immunization and promptly yield COVID-19 vaccine prototypes for further translational development.

**Significance:** The ongoing COVID-19 global pandemic has accounted for over 2.5 million deaths and an unprecedented impact on the health of mankind worldwide. Over the past several months, while a few COVID-19 vaccines have received Emergency Use Authorization and are currently being administered to the entire human population, the demand for prompt global immunization has created enormous logistical challenges--including but not limited to supply, access, and distribution--that justify and reinforce the research for additional strategic alternatives. Phage are viruses that only infect bacteria and have been safely administered to humans as antibiotics for decades. As experimental proof-of-concept, we demonstrated that aerosol pulmonary vaccination with lung-targeted phage particles that display short epitopes of the S protein on the capsid as well as preclinical vaccination with targeted AAVP particles carrying the S protein gene elicit a systemic and specific immune response against SARS-CoV-2 in immunocompetent mice. Given that targeted phage- and AAVP-based viral particles are sturdy yet simple to genetically engineer, cost-effective for rapid large-scale production in clinical grade, and relatively stable at room temperature, such unique attributes might perhaps become additional tools towards COVID-19 vaccine design and development for immediate and future unmet needs.

## Introduction

Since the beginning of 2021, the World Health Organization has estimated that over 2.5 million deaths in 192 countries/territories have been caused by complications of coronavirus disease 2019 (COVID-19). This unprecedented pandemic has prompted a worldwide collaborative effort to develop antiviral therapies and vaccines to control global spread. Severe acute respiratory syndrome coronavirus 2 (SARS-CoV-2) is the third zoonotic coronavirus to infect humans in less than 20 years (1, 2). Previous coronavirus epidemics, such as SARS-CoV and Middle East Respiratory Syndrome (MERS), foreshadowed the risk of emergent outbreaks and the imminent need for novel and versatile technologies for rapid manufacturing and large-scale distribution of vaccines and therapies as an emergency countermeasure.

SARS-CoV-2 is a single-stranded enveloped RNA virus with four main structural proteins. The spike (S) protein mediates both host cell recognition and membrane fusion and is pivotal for viral entry. The S protein, comprised of the S1 and S2 subunits, is displayed as a trimer on the surface of the viral particle (3). Within the S1 subunit, the receptor-binding domain (RBD) adopts an open conformation that can interact with the angiotensin-converting enzyme 2 (ACE2) receptor on the host cell membrane. Upon binding, the S1 subunit is cleaved and subsequent conformational changes in the S2 subunit trigger the formation of a six-helical bundle comprised of heptapeptide repeat sequence 1 (HR1) and heptapeptide repeat sequence 2 (HR2), followed by the insertion of the fusion peptide (FP) into the host cell membrane. Given the importance of the S protein for the entry of SARS-CoV-2 in the host cells, it has served as the main target for vaccines and therapeutic antibodies. Thus, the identification and understanding of structurally-defined S protein epitopes with potential neutralizing capabilities are crucial in the design of effective and robust vaccines and antibody cocktail therapies (4, 5).

Current vaccine platforms against COVID-19 are classified into several broad categories that include nucleic acid-based vaccines (mRNA and DNA), viral vector vaccines (e.g., adenovirus), inactivated or live attenuated viral vaccines, or recombinant protein or peptide vaccines. Some have been granted Emergency Use Authorization by various regulatory agencies, including two mRNA vaccines (Pfizer-BioNtech and Moderna), three non-replicating adenovirus vaccines (Oxford/AstraZeneca, Sputnik V, and Johnson & Johnson), and an inactivated virus vaccine (CoronaVac), while many others are pending approval or undergoing clinical trials. In addition to such extraordinary global efforts, further exploration and development of vaccines that are amenable to temperature fluctuations, rapid large-scale production and distribution, and conferral of long-term immunological protection in the face of existing and emerging variant strains remain an unmet need (6, 7).

Phage particles have been used in clinical settings for nearly a century, and represent an inexpensive, and versatile tool for large-scale immunization. Phage are viruses that naturally infect bacteria; during the pre-antibiotic era, humans received phage to neutralize systemic bacterial infections without significant adverse effects (8, 9). Recently, engineered phage particles have been leveraged in different applications, particularly in vaccine development because they are: (i) easy to genetically engineer and produce in large scale in *Escherichia coli*; (ii) strong immunogens capable of stimulating both cellular and humoral immunity (10-12); (iii) highly stable under harsh environmental conditions (pH and temperature). Such features facilitate administration, storage, and transport (10); (iv) importantly, they are considered safe for administration to humans (9, 13-15). We and others have shown that the addition of peptide targeting sequences to the minor coat protein pIII and immunogenic peptide sequences on the recombinant pVIII coat protein (rpVIII) confer tropism of phage to specific eukaryotic receptors in normal or diseased tissue or to organelles (16-21). Pertinent to vaccine development, we recently demonstrated that aerosolized phage particles targeting the lung and subsequently undergoing receptor-mediated transport to the systemic circulation are safe and elicit specific and sustained local and systemic immune responses in mice and non-human primates (22). Further genome engineering was leveraged to create a hybrid adeno-associated virus/phage vector (20), which has been extensively validated for ligand-directed gene delivery and represents a viable alternative for nucleotide-based vaccines (mRNA or DNA). Several favorable features were built into the vector design, including: (i) adjuvant properties of phage particles, (ii) ligand-directed system for receptor-mediated internalization, (iii) well-characterized fate of the genome (concatemerization and integration of intact genomes), and (iv) ability to avoid neutralization after an immune response against phage, as proven in repeat-dose studies using pet dogs (23) and several animal models (20, 23-28).

We, therefore, explored the inherent biological and genetic properties of phage particles to produce novel vaccine candidates against COVID-19. First, guided by structure-based antigen design, we selected S protein epitopes that were genetically incorporated into the recombinant major capsid coat protein pVIII (rpVIII) of the f88-4 phage genome. The peptide sequence CAKSMGDIVC, which was recently described for its unique ability to target lung epithelium cells and induce the transport of phage particles into the systemic circulation (22), was then cloned into phage minor coat protein pIII of the fUSE55 phage genome, creating a dual-display system for efficient delivery <u>and</u> immunization against the SARS-CoV-2 S protein. This approach allows us to use the lung as the main route of vaccination, which has remarkable advantages over conventional routes. Aerosol administration is needle-free and requires minimal use of specialized medical staff; it is not subject to first-pass metabolism and is considered the most effective route for inducing local immune protection against airborne pathogens (29-32). In a second approach, we used the AAVP vector (20), which contains a eukaryotic expression cassette flanked by AAV inverted terminal repeat (ITR) sequences within the phage genome, for the delivery and transduction of the full-length S protein gene (AAVP *S*) in host cells. This approach leverages the same rationale that serves as the basis for the activity of existing vaccines (mRNA or adenovirus) that trigger a specific humoral response against the full-length S protein. Integration of large DNA antigen sequences flanking the AAV transgene of the AAVP confers stability to the construct upon transduction and enhanced expression efficiency. Both phage- and AAVP- based vaccines elicited systemic S protein-specific humoral response in mice with no evidence of adverse effects, indicating that both technologies hold promise in vaccine development.

## Results

### Development of novel phage- and AAVP-based vaccine platforms against SARS-CoV-2

We pursued two different strategies for immunization: (i) phage-based vaccine candidates displaying various S protein epitopes, and (ii) an AAVP-based vaccine candidate that utilizes the entire SARS-CoV-2 S protein (Fig. 1). In both approaches, we incorporated a ligand peptide along with the viral antigens in the phage or AAVP to target specific cell surface receptors and facilitate the immune response.

**Fig 1.**
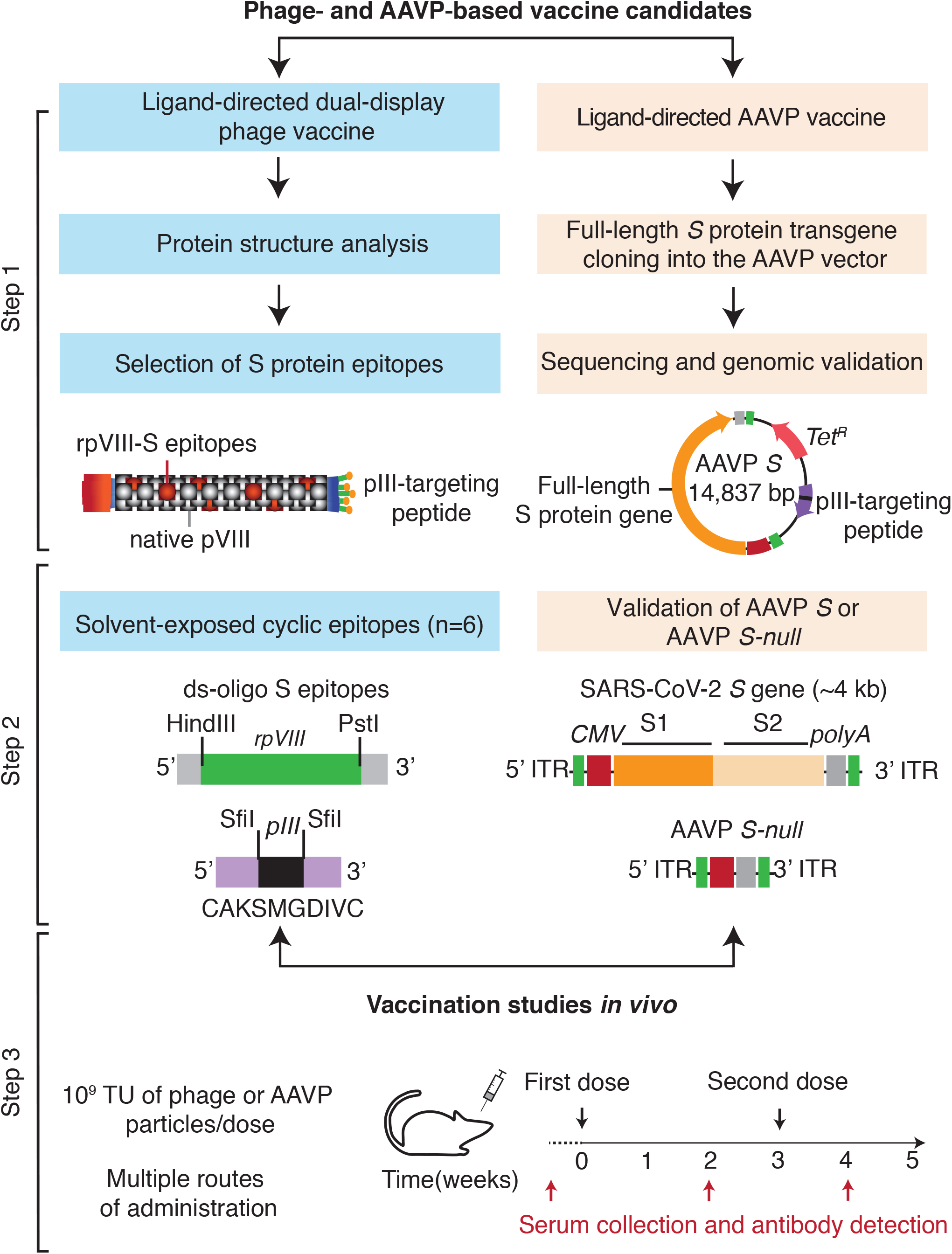
Schematic representation of the phage- and AAVP-based vaccine candidates. The scheme represents the approach used for the conception, design, and application of two strategies of immunization against SARS-CoV-2 S protein using phage particles. Step 1: Structural analysis, selection of structurally-defined epitopes, and cloning steps for the generation of dual-display phage particles and AAVP encoding the full-length S protein. Step 2: Molecular engineering of single- and dual-display phage particles, and AAVP *S* constructs. Step 3: Functional validation and vaccination studies *in vivo* in mice.

In the capsid engineering system, we genetically engineered phage to display immunologically relevant S protein epitopes (see below) on the highly exposed rpVIII protein of the phage capsid using the f88-4 vector (Fig. 1, Step 1) (18, 19). To enable tissue-specific targeting of these phage particles, we also subcloned the coding sequence of the novel CAKSMGDIVC peptide ligand into the pIII gene of the fUSE55 vector, yielding a dual-display phage (Fig. 1, Step 2). The CAKSMGDIVC ligand binds to α3β1 integrins and mediates the transport of phage particles across the lung epithelium to the systemic circulation where they elicit strong and sustained pulmonary and systemic humoral responses against antigens displayed on the phage capsid (22). As a control, we used the untargeted parental phage particles (insertless phage) that display the native pVIII and pIII proteins.

For our second strategy, based on gene delivery, we inserted an expression cassette containing the full-length S protein transgene and the human *CMV* promoter in *cis* conformation within the 5’ and 3’ ITRs in the AAVP genome for gene delivery and transduction in host cells (Fig. 1, Step 2). As a control, we used the targeted AAVP empty vector (termed “AAVP *S-null*”). Targeting was established by display of another integrin-binding peptide, ACDCRGDCFCG (RGD4C) on the pIII protein, which has high affinity for α_v_ integrins (17, 33), which are highly expressed in leukocytes trafficking to draining lymph nodes and areas of inflammation (34). The RGD motif (arginine-glycine-aspartate) facilitates particle uptake by dendritic cells and enhances the immunogenicity of peptide antigens, DNA vaccines, and adenovirus vectors (35-37).

Both approaches allow the rapid “swapping” of targeting motifs, epitopes, and/or gene coding sequences, thus providing rapid and powerful design flexibility to produce a variety of vaccines and overcome potential limitations in protein conformation of structure-based epitopes. The dual-display phage particles, the RGD4C-AAVP *S* particles, and corresponding controls were tested *in vivo* in mice to assess different routes of administration, and to evaluate the induced antigen-specific humoral response by ELISA (Fig. 1, Step 3). The overall vaccination schedule included at least two administered doses of 10^9^ transducing units (TU) of phage or AAVP particles with an interval of 1–2 weeks.

### Identification and selection of epitopes for dual-display phage-based vaccine

To identify relevant epitopes for phage capsid manipulation, *in silico* analysis of the experimentally-determined viral S protein structure of the Wuhan-Hu-1 strain (GenBank Accession number: NC_045512.2) was performed. We prioritized solvent-exposed amino acid stretches with flanking cysteine residues and cyclic conformation, because these amino acid sequences are more likely to recapitulate the composition of endogenous epitopes and thereby increase the likelihood of antigen recognition and processing by the host immune system. Other epitopes were also considered following structure-guided predictions, even in the absence of flanking cysteine residues. Also, given that phage particles are produced in prokaryotic host cells, we prioritized epitopes lacking sites expected to undergo post-translational modification.

We selected six S protein epitopes, which are accessible in both the closed and open states of the S protein. At least five of these epitopes have since been shown to be fully or partially immunogenic (Fig. S1 and Table S1) (38-45). The six epitopes range in length from 9 to 26 amino acids (aa). Four occur within the S1 subunit and two are found in the S2 subunit (Fig. 2A). Three of the S1 epitopes are located in the receptor-binding domain (RBD): epitope 1 (aa 336–361), epitope 2 (aa 379–391), and epitope 3 (aa 480–488). The remaining epitope derived from the S1 subunit, epitope 4 (aa 662–671), is located near the cleavage site between the S1 and S2 subunits. Epitopes within the S2 subunit, epitope 5 (aa 738–760) and epitope 6 (aa 1032–1043), are located near FP (aa 788–806) and HR1 (aa 912–984), respectively. Most of the selected epitopes are cyclic due to the presence of flanking cysteine residues, except epitope 2 (aa 379–391), which adopts a loop-like conformation despite the absence of disulfide bridges (Fig. 2B).

**Fig 2.**
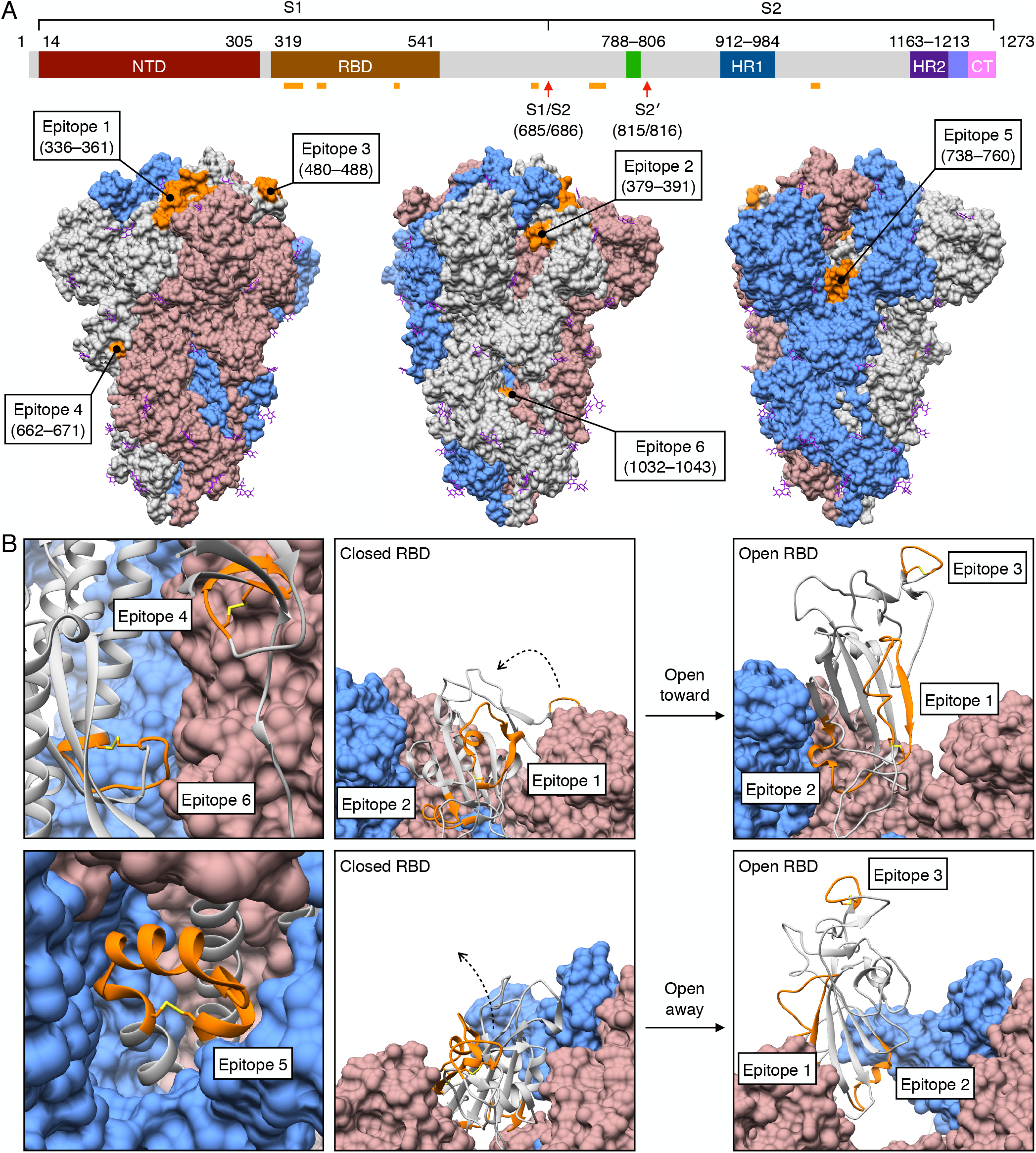
Identification of structural epitopes on S protein trimer. (A) Six epitopes (orange) spanning the SARS-CoV-2 S protein were selected for display on rpVIII protein. Four epitopes are located within the S1 subunit: epitope 1 (aa 336–361), epitope 2 (aa 379–391), epitope 3 (aa 480–488), epitope 4 (aa 662–671); and two within the S2 subunit: epitope 5 (aa 738–760), and epitope 6 (aa 1032–1043). These epitopes are solvent-exposed in the surface representation of the predominantly closed-state conformation of the S protein trimer (gray, cornflower blue, and rosy brown) (PDB ID: 6ZP0) (60). Only epitope 1 (aa 336–361) contains a site for glycosylation (at N343) (purple). (B) All of the epitopes (orange) maintain a cyclic conformation in the ribbon representation of a S protein protomer (gray); disulfide bridges (yellow) are present between the flanking cysteine residues of all epitopes except on epitope 2 (aa 379–391). The open-state conformation of the S protein trimer with one RBD erect displays a change in orientation of epitopes 1, 2, and 3, though all remain solvent-exposed (PDB ID: 6ZGG) (60).

Many studies have demonstrated that the S protein is highly glycosylated, and some glycosylation sites have been reported to alter the infectivity of variants and facilitate evasion of the host immune response (4, 7). Considering that we selected epitopes based on conformation, we identified that epitope 1 contains a glycosylation site on residue N343. Notably, this site seems to be important in viral infectivity in which the glycosylation deletion N343Q drastically reduces the D614G variant infectivity (7). In our system, however, we expect that the lack of N-glycosylation will not produce a significant structural divergence in the epitope conformation when displayed on the phage capsid, as similarly observed with other SARS-CoV-2 strains (46). Because the glycosylation site of epitope 1 is located in its N-terminus and unlikely to interrupt antibody recognition of the remaining structure, we sought to investigate the efficacy of this additional epitope.

### Epitope conformational analysis by molecular dynamics

To examine whether the predicted conformation of each epitope on display on the phage capsid would recapitulate its features on the S protein, we performed *in silico* conformational analysis using microsecond-scale all-atom explicit-solvent simulations of each epitope. We hypothesized that epitope conformation is a pre-requisite for successful induction of the immune response against the native S protein. Notably, we found that epitope 4 has the lowest spatial root-mean-square deviation (RMSD) amongst the four S1 subunit epitopes. While its RMSD fluctuates between 2 Å and 4 Å (Fig. 3A), epitope 4 has the most frequent low values (RMSD∼2Å), which may explain its near-native conformation when compared to the structure of the S protein (Fig. 3B) (47). As for the other selected epitopes of the S1 subunit (epitopes 1, 2, and 3) and the S2 subunit (epitopes 5 and 6), RMSD values are substantially higher (4–10 Å) (Fig. 3C, D), after an initial relaxation of each system (∼100 nanoseconds). These differential RMSD trends suggest that epitope 4 is most likely to resemble the native conformation and generate IgG antibodies with high cross-reactivity with the S protein. If true, epitope 4 represents a promising candidate for display on phage capsid that would elicit an immune response towards an exposed region of the S protein. It would also demonstrate that molecular dynamics simulations can serve as a useful tool for prioritizing epitopes with higher likelihood of recapitulating native conformation when selecting epitope candidates for use in phage-based vaccines.

**Fig 3.**
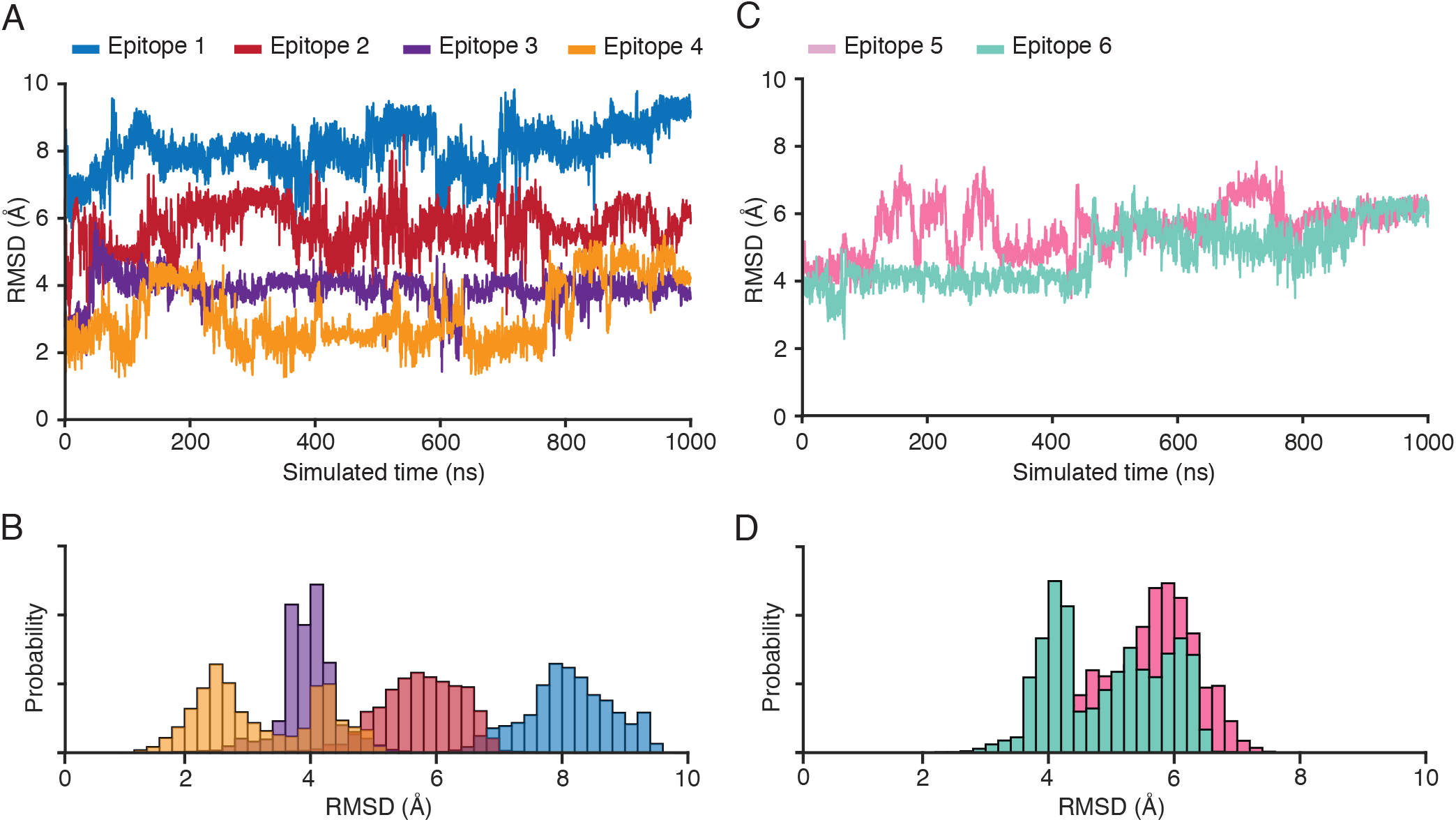
Explicit-solvent simulations reveal the near-native conformation of epitope 4 relative to the full-length S protein. (A) RMSD from the S protein conformation for each epitope within the S1 subunit: epitope 1 (aa 336–361), epitope 2 (aa 379–391), epitope 3 (aa 480–488) and epitope 4 (aa 662–671). Epitope 4 frequently samples low values (∼2Å), while the other epitopes undergo substantial conformational rearrangements (RMSD<4Å). (B) Probability as a function of RMSD shows that epitope 4 is most likely to sample conformations that are similar (RMSD ∼2.5Å) to the S protein conformation. (C) RMSD as a function of time for the epitopes within the S2 subunit: epitope 5 (aa 738–760) and epitope 6 (aa 1032–1043). Both epitopes rapidly adopt and maintain large RMSD values (D).

### Characterization of structurally-defined S epitopes for immunogenicity in mice

To evaluate the immunological potential of each of S epitope and select promising candidate(s) for phage-based vaccine development, we tested their ability to induce an immune response in mice. Six f88-4 phage were produced, each exhibiting one of the six S protein epitopes fused into the rpVIII protein on ∼300 copies per phage particle (termed “single-display phage”) (Fig. S2).

Immunogenicity of the epitopes expressed on the rpVIII protein was assessed in mouse serum (Swiss Webster or BALB/c) obtained after the first dose (prime) and the second dose (boost) administered subcutaneously and compared against the control insertless phage. Antigen-specific IgG titers were quantified by ELISA using an immobilized recombinant S protein (aa 16–1213) for capture. Epitope 4 from the S1 subunit induced high levels of S protein-specific IgG antibodies, and booster immunizations further increased antibody levels (Fig. 4A). None of the other five phage constructs generated S protein-specific IgG antibodies above the background levels observed for mice immunized with the control insertless phage. These results indicate that epitope 4 is the most immunogenic among the selected epitopes, suggesting that epitope display of the native conformation supports development of specific immune response as predicted by the *in silico* analysis.

**Fig 4.**
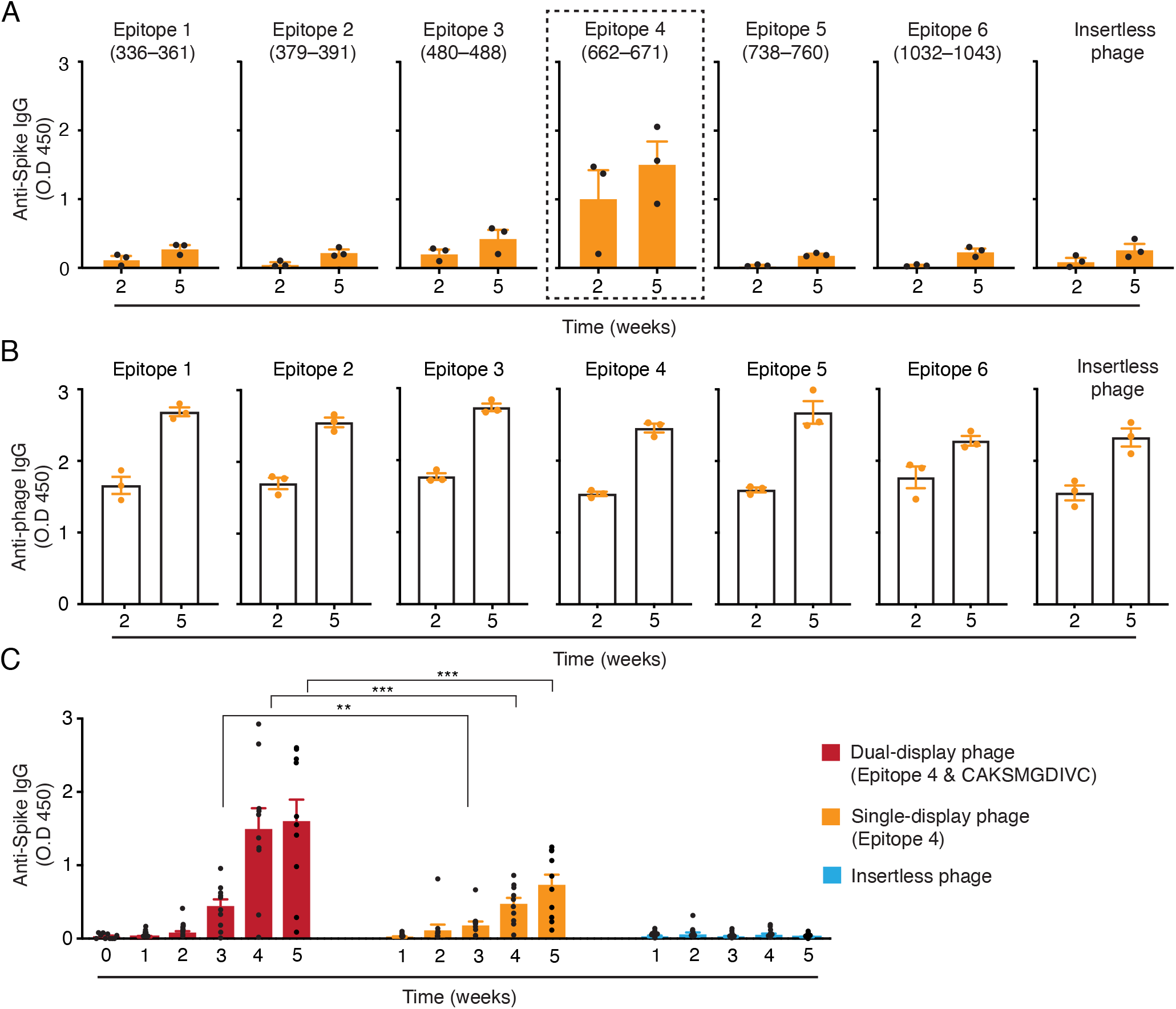
Immunogenicity of S protein epitopes on single-display phage particles. Five-week-old female Swiss Webster mice were immunized via subcutaneous injection with single-display phage constructs containing each of the six different epitopes expressed on rpVIII protein or the control insertless phage. Animals received a boost injection three weeks after the first administration. (A) S protein-specific IgG antibodies and (B) phage-specific IgG antibodies were evaluated in sera of mice after two- and five-weeks by ELISA (n=3 mice per group). (C) Five-week-old female BALB/c mice were immunized via intratracheal administration with the epitope 4/CAKSMGDIVC dual-display phage particles, epitope 4 single-display phage particles, or the control insertless phage. Animals received a boost three weeks after the first administration. S protein-specific IgG antibodies were weekly evaluated by ELISA (n=10 mice per group). Data represent ± SEM (* *P* < 0.05, ** *P* < 0.01, *** *P* < 0.001).

Given the well-documented inherent immunogenicity of native filamentous phage, we also investigated the levels of phage-specific IgG antibodies in the sera of these mice. Notably, all single-display phage constructs generated high titers of phage-specific IgG antibodies, which were markedly increased after the second dose independent of the presence or absence of the S protein epitopes displayed on the rpVIII protein (Fig. 4B). It is remarkable that generation of phage-specific IgG antibodies does not appear to compromise the S protein-specific humoral response. A clear distinction was observed between epitope 4 and the other phage particles, including the control insertless phage. Epitope 4, therefore, was selected as the lead candidate for testing a dual-display phage-based vaccine bearing a targeting moiety.

### Dual-display phage construct for pulmonary vaccination

To generate a dual-display phage-based vaccine, we optimized a simple two-step cloning strategy that allows rapid exchange of epitopes and/or targeting peptide ligands in the phage genome. This methodology has the potential to rapidly mitigate the shortcomings in the neutralization due to possible mutations and/or direct the phage to target cells or tissue to improve the immune response. Pulmonary vaccination has been shown to be the most efficient route to generate mucosal and systemic immunity against airborne pathogens (22, 29-31) and significantly increase immunological protection of non-human primates infected with SARS-CoV-2 (30, 32). Therefore, we prioritized the peptide CAKSMGDIVC for lung targeting and aerosol delivery (22) and generated dual-display phage particles that simultaneously express epitope 4 on rpVIII protein (∼300 copies) and the peptide CAKSMGDIVC on pIII protein (3-5 copies).

To determine the immunogenic properties of the epitope 4/CAKSMGDIVC dual-display phage particles, groups of five-week-old BALB/c female mice were dosed intratracheally. Cohorts of mice (n=10 per group) received two doses of 10^9^ TU of the epitope 4/CAKSMGDIVC dual-display phage, or the epitope 4 single-display phage, or the control insertless phage in 3-week intervals. The presence and levels of S protein-specific IgG antibodies were evaluated in serum samples collected weekly by ELISA. Titers of S protein-specific IgG antibodies were higher in mice immunized with the epitope 4/CAKSMGDIVC dual-display phage particles compared to the control insertless phage, especially after three weeks of the first dose, and with a substantial increase after the second dose (weeks 4 and 5) (Fig. 4C). Epitope 4 single-display phage particles also induced systemic S protein-specific IgG antibodies but do so at levels lower than the dual-display phage particles. These results indicate that the addition of the CAKSMGDIVC peptide ligand mediates the transport of the dual-display phage from the pulmonary tree to the systemic circulation, thereby increasing the immunogenicity.

Together, our findings demonstrate that epitope 4 induces a robust S protein-specific humoral response when displayed on the rpVIII protein as a single display entity (single-display phage particles) and, when combined with the lung transport peptide CAKSMGDIVC, immunogenicity is enhanced. These results provide encouraging evidence that epitope 4/CAKSMGDIVC dual-display phage particles are suitable candidates for pulmonary vaccination against SARS-CoV-2 and pave the way for further validation of phage-based cocktails as vaccines that cover a broad spectrum of antigenic sites that elicit the production of antibodies with well-established and/or new neutralizing capabilities.

### A novel AAVP-based vaccine for efficient gene delivery and humoral response against the viral S protein

As a parallel approach to the dual-display phage design, we adapted a long-standing and well validated targeted AAVP gene therapy platform (20) to deliver the S protein gene (Wuhan-Hu-1 strain, GenBank Accession Number: NC_045512.2) (termed “AAVP *S*”). AAVP *S* displays the ACDCRGDCFCG (RGD4C) targeting peptide ligand on the pIII protein, allowing it to bind α_v_ integrins. These integrins are known to regulate trafficking of lymphocytes and antigen-presenting cells (i.e., dendritic cells) into secondary lymphoid organs (37). For the control, we generated a RGD4C-AAVP empty vector that is identical to RGD4C-AAVP *S* but does not contain the S protein gene (RGD4C-AAVP *S-null*) (Fig. 5A, B).

**Fig 5.**
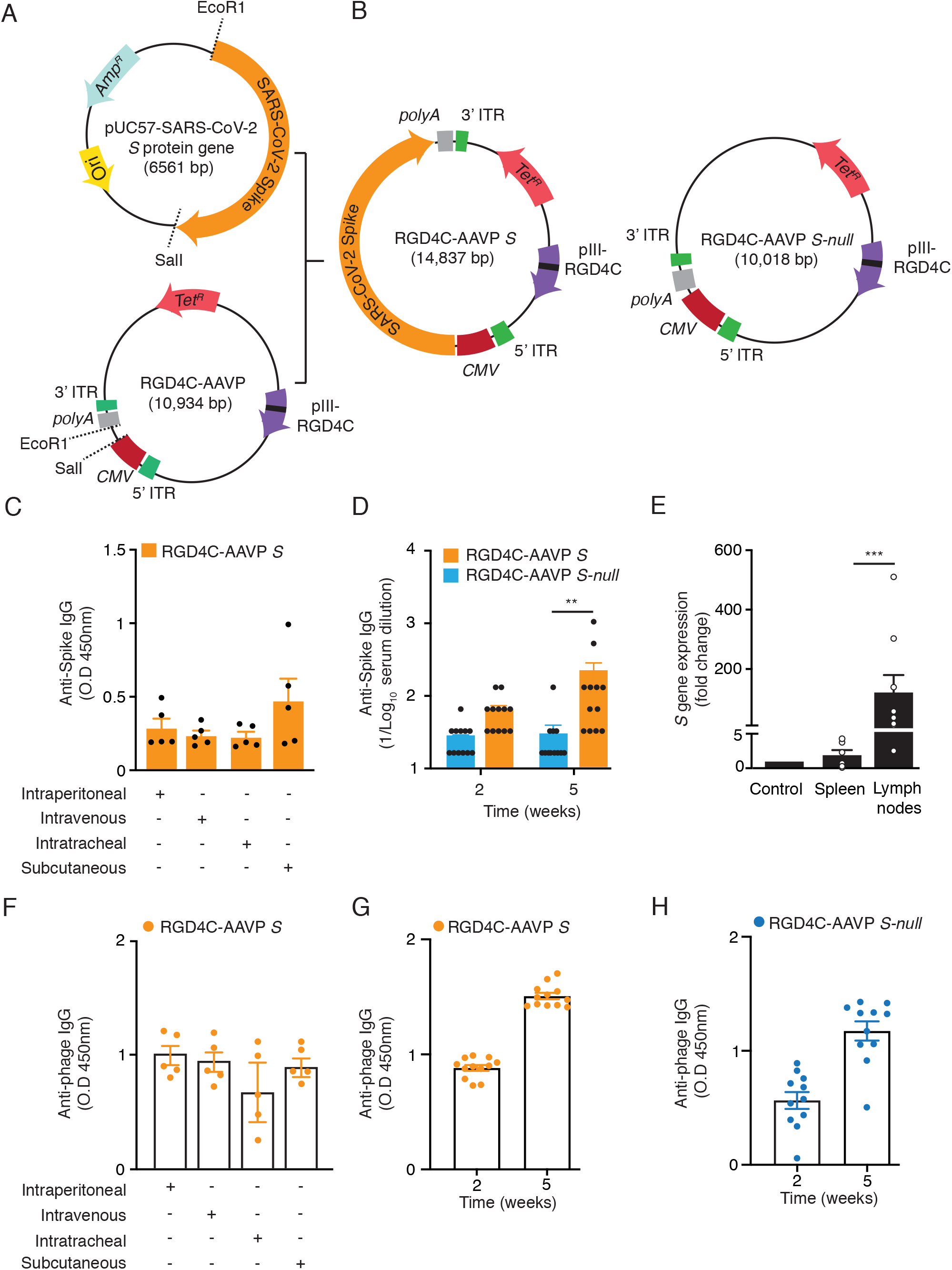
Immunogenicity of the RGD4C-AAVP *S* in mice. Schematic representation of the AAVP-based vaccine candidate. (A) The modified, synthetic SARS-CoV-2 *S* protein gene was excised from the pUC57 and cloned into the RGD4C-AAVP -TNFΔEcoRI829. Expression of the *S* protein transgene cassette is driven by the cytomegalovirus (*CMV*) promoter and flanked by AAV ITRs. (B) Schematic representation of the RGD4C-AAVP *S* and the control RGD4C-AAVP empty vector (RGD4C AAVP *S-null*). (C) S protein-specific IgG antibody response in the sera of mice immunized with RGD4C-AAVP *S* via different routes of administration (n=5 mice per group) by ELISA. (D) S protein-specific IgG antibodies in sera of mice weekly immunized with RGD4C-AAVP *S* or the control RGD4C-AAVP *S-null* (n=12 mice per group) via subcutaneous administration. Data represent ± SEM (** *P* < 0.01). (E) Tissue-specific expression of the *S* protein transgene in mice immunized with AAVP *S* five weeks after the first immunization. Data represent ± SEM (*** *P*< 0.001). (F) Phage-specific IgG antibody response in the sera of mice immunized with RGD4C-AAVP *S* via different routes of administration (n=5 mice per group). Phage-specific IgG antibody response in mice immunized with RGD4C-AAVP *S* (G) or RGD4C-AAVP *S-null* (H) at two and five weeks after the first immunization. Phage-specific IgG antibody response was evaluated by ELISA in 96-well plates coated with 10^10^ AAVP particles per well. Tet^R^, tetracycline resistance gene. Amp^R^, ampicillin resistance gene. Ori origin of replication.

To assess the immunogenicity of RGD4C-AAVP *S*, we first immunized five-week-old female outbred Swiss Webster mice. Cohorts of mice (n=5) received 10^9^ TU of targeted RGD4C-AAVP *S via* intraperitoneal (group 1), intravenous (group 2), intratracheal (group 3), or subcutaneous (group 4) routes. The S protein-specific IgG antibody response was evaluated 14 days later by ELISA (Fig. 5C). Baseline sera were used as controls. Administration of RGD4C-AAVP *S* particles elicited serum IgG responses against the S protein in all experimental groups, whereas mice immunized *via* subcutaneous injection had higher serum IgG titers than the other groups. Thus, we elected to administer RGD4C-AAVP *S* subcutaneously for subsequent *in vivo* assays.

We tested our immunization regimen in five-week-old female inbred BALB/c mice. Cohorts of mice (n=12 per group) received weekly subcutaneous doses of 10^9^ TU of RGD4C-AAVP *S* or the control, RGD4C-AAVP *S-null*. Higher titers of S protein-specific IgG antibodies were observed in mice vaccinated with RGD4C-AAVP *S* relative to the baseline sera and RGD4C-AAVP *S-null*, particularly five weeks after the first dose, confirming that RGD4C-AAVP *S* is a suitable vector for transgene delivery and elicits a systemic humoral immune response (Fig. 5D).

To gain insight into the S protein transgene expression mediated by RGD4C-AAVP *S*, we investigated the fate of the phage genome, namely, the sites in which transduced cells were detected in mouse tissues, including main regional lymph nodes (axillary, inguinal, mesenteric, and mediastinal) 4 weeks after the first dose. Transgene expression at varying levels was detected mainly in the draining lymph nodes. Skeletal muscle and spleen were used as control organs and show only background levels of transgene expression (Fig. 5E). As expected, transcripts for the S protein were not detected in mice immunized with RGD4C-AAVP *S-null*. These results confirm that gene delivery by AAVP and the expression of the S protein in the draining lymph nodes trigger a systemic S protein-specific humoral response. Moreover, the data recapitulate the well-established attribute of AAVP particles in preventing off-target effects, even upon clearance *via* the reticulum-endothelial system (RES), sparing non-targeted or distal tissues, while a strong promoter drives the expression of the transgene in the transduced cells (20, 23-28). This finding is particularly important for evaluating potential adverse effects in novel, candidate AAVP-based vaccines, since off-site transduction-associated toxicities have been reported in toxicological studies of adenovirus vaccines (48, 49). Therefore, the extensive body of data generated to date with AAVP in cancer gene therapies (20, 23-28) can help accelerate future clinical development of AAVP vaccine candidates, potentially in a highly cost-effective and efficient manner.

Finally, we examined the antibody response against phage in mice vaccinated with RGD4C-AAVP *S* or RGD4C-AAVP *S-null* phage. We observed a strong and sustained phage-specific IgG antibody response upon administration of RGD4C-AAVP *S* by all routes of administration (Fig. 5F) and the response increased after boost injections in both RGD4C-AAVP S (Fig. 5G) and RGD4C-AAVP *S-null* (Fig. 5G), indicating that AAVP are strong immunogens and likely serve as adjuvants for AAVP-based vaccines. Together, these results indicate that targeted AAVP *S* is an efficient tool for delivery and expression of viral proteins and can induce an immune response against the S protein.

## Discussion

In this study we designed, generated, and evaluated the translational potential of phage- and AAVP-based vaccine candidates using capsid epitope display and gene delivery as strategies for immunization against SARS-CoV-2. We demonstrated that both systems can be successfully used to induce an antigen-specific humoral response against the S protein and therefore represent valid candidate approaches for vaccine development. Notably, as part of ongoing work, we have optimized Good Manufacturing Practice (GMP) generation, production, and purification of engineered phage particles so that manufacturing could be accomplished on an industrial scale for rapid commercialization.

One of the main challenges associated with vaccines deployed under Emergency Use Authorization (EUA) is to predict the potency and duration of the protective immune response against host cell-engaging epitopes on the S protein, particularly in the face of new genetic variants. In principle, focusing on structural antigen mapping and immunodominant B-and T-cell epitopes that trigger immune responses associated with both potent neutralizing activity and lower transmissibility would lead to long-term protection. As such, many studies have aimed to predict an/or map epitopes from B- and T-cells derived from the SARS-CoV-2 S protein and other structural proteins (50). In this study, we selected six exposed regions of the S protein with specific structural constraints for display on the phage capsid and to increase the likelihood of antigen recognition and processing by the host immune system. We found that epitope 4 (aa 662–671) triggered a strong and specific systemic humoral response against the S protein, presumably by recapitulating the near-native conformation of the epitope when expressed on the rpVIII protein as predicted by molecular dynamics analyses. Thus, the combinatorial approach of selecting regions of an antigen based on conformational constraints and evaluating their structural dynamics *in silico* can be used to identify epitopes likely to replicate the natural immune response to an infection. Our findings suggest that antigen-engineering strategies, such as the vaccine candidate for the Zika virus (51), have the potential to generate vaccines with high efficacy in inducing a broadly functional repertoire of neutralizing antibodies and cell-mediated immune responses.

To support the translational applications of phage-based vaccination, we designed a protocol for immunization in mice as a proof of principle towards pulmonary vaccination against SARS-CoV-2. The design of dual-display phage particles relies on a clever, yet simple bioengineering exercise: the simultaneous display of both epitope 4 on the pVIII protein and the CAKSMGDIVC targeting ligand on the pIII protein. Because CAKSMGDIVC mediates selective targeting and transport of phage particles to the systemic circulation (22), we showed that an aerosol strategy of immunization may confer biologically significant advantages over conventional routes of immunization. First, unlike subcutaneous or intramuscular injections, inhalation is needle-free and minimally invasive, eliminating the requirement for specialized medical staff, and leading to the potential for self-administration for isolated and/or vulnerable populations sheltering in place, particularly the elderly and the immunocompromised. Moreover, rapid access to the upper and lower respiratory tract leads to a reduction in viral shedding, which results in lower transmission (52-54).

Antigen exposure to the lung surface, which is lined by the highly vascularized pulmonary epithelium, is a unique feature that is known to induce a local immune response that decreases infection and transmission of airborne pathogens. Emerging studies of intranasal or intratracheal immunizations have already proven successful protection against SARS-CoV-2 in mice and non-human primates (32, 54).

We also uncovered the potential value of using AAV-Phage particles to deliver the S protein gene as an additional alternative SARS-CoV-2 vaccine candidate. In the last decade, AAVP technology has proven to be a modular platform that can be appropriately tailored to image and treat a variety of human solid tumors in mouse models and spontaneous tumors in pet dogs (20, 23-28). These attributes make AAVP a unique platform for gene delivery. Indeed, we showed that administration of RGD4C-AAVP *S* particles elicits an antibody response in mice against the encoded S protein. Because our prototype AAVP *S* vaccine is targeted with the integrin-binding peptide (RGD4C), which has a high affinity binding for α_v_ integrins, this approach may facilitate targeting of inflammatory cells trafficking to the lymph nodes where gene expression and antigen presentation occur. The identification of a RGD motif within the RBD domain of the S protein suggests that integrins may act as co-receptors or an alternate path for coronavirus entry (55). Therefore, it is plausible that different functional ligands for tissue-specific transgene expression within lymph nodes (12), lymphatic vessels (56), or lung epithelial cells (e.g., CAKSMGDIVC) which has proven to be highly efficient in inducing local and systemic immune response upon pulmonary delivery (22), may enhance the efficacy and broad administration of AAVP-based vaccines.

In conclusion, here we bring forward and validate the first proof-of-concept for the design, structure-function relationship, and translation of both phage- and AAVP-based vaccines against SARS-CoV-2. A targeted pulmonary vaccination strategy with a simplified supply chain may also be applicable for potential utility against other viruses.

## Materials and Methods

### Animals

Four-to-six-week-old Swiss Webster and BALB/c mice were purchased from the Jackson Laboratory (Sacramento, CA) and were housed in specific pathogen and opportunist free (SOPF) rooms with controlled temperature (20 ± 2°C), humidity (50 ± 10%), light cycle (light, 7:00-19:00; dark, 19:00 – 7:00), and access to food and water *ad libitum* at the research animal facilities of the Rutgers Cancer Institute of New Jersey. Littermates were randomly assigned to experimental groups. The Institutional Animal Care and Use Committee (IACUC) from the Rutgers Cancer Institute of New Jersey approved all animal experiments.

### Structural Analysis of S Protein for Epitope Selection

The structure of SARS-CoV-2 S protein (PDB IDs: 6VXX, 6VYB) (4) was analyzed using UCSF Chimera software (57) for selection of epitopes to display on rpVIII protein. Though epitope 3 (aa 480–488) was not resolved in these early structures, the flanking cysteine residues of this region were predicted to form a disulfide bridge (58). This has been confirmed experimentally with since-determined structures (e.g., PDB IDs: 6ZP0, 6ZGG) (59, 60).

### Molecular Dynamics Simulations

All-atom explicit-solvent simulations of the epitope sequences were performed with the GROMACS 2020 software package (61, 62). The initial configuration for each epitope was taken from a cryo-EM structure of the full-length S protein (PDB ID: 6XR8) (63). Each epitope was solvated using the TIP3P water model (64), where the box size was defined to have a 10 Å buffer between the edge of the box and the epitope. Depending on the charge of each molecule, neutralization with either Cl^-^ or Na^+^ ions was applied. The AMBER99SB-ILDN protein force field (65) was used for all simulations. After steepest-descent energy minimization, each system was equilibrated at 270 K using the NVT ensemble for 5 nanoseconds (ns), followed by the NPT ensemble for 5 ns, while position restraints were imposed on all heavy atoms (1000 kJ/nm^2^). Restraints were then removed and steepest-descent minimization was performed, followed by NVT and NPT equilibration simulations (5 ns each, at 270 K). All production runs were performed in the NPT ensemble using the Nose-Hoover thermostat (61, 66) at 310 K and the Parrinello-Rahman barostat (67) set at 1 bar. Each production simulation was performed for a minimum of 1 microsecond (ms) (21 ms, aggregate simulated time). To ensure reproducibility of the results, a second set of simulations were equilibrated using identical protocols, except that the temperature during equilibration was 310 K. The overall dynamics were not sensitive to the equilibration temperature.

### Generation of S Protein Epitopes Single-Display Phage Particles

To generate single-display phage constructs, we used the fd-tet-derived vector f88-4 containing a recombinant gene VIII (GenBank Accession Number: AF218363.1). Single colonies were selected on Luria-Bertani (LB) agar plates with tetracycline (40 μg/mL) and cultured overnight (O.N.). Each plasmid DNA was isolated by plasmid purification (Qiagen). Next, annealed oligonucleotides encoding for each of the six selected epitopes: epitope 1: fwd 5’AGCTTTGCCTGTCCGTTCGGCGAAGTGTTCAACGCGACCCGCTTCGCGAGCGTGTATGCGTGGAACCGCAAACGCATCAGCAACTGTCCTGCA 3’, rev 5’ GGACAGTTGCTGATGCGTTTGCGGTTCCACGCATACACGCTCGCGAAGCGGGTCGCGTTGAACACTTCGCCGAACGGACAGGCAA 3’; epitope 2: fwd 5’ AGCTTTGCCTGTTATGGCGTGAGCCCGACCAAACTGAACGATCTGTGTCCTGCA 3’, rev 5’ GGACACAGATCGTTCAGTTTGGTCGGGCTCACGCCATAACAGGCAA 3’; epitope 3: 5’ AGCTTTGCCTGTAACGGCGTGGAAGGCTTCAACTGTCCTGCA 3’, rev 5’ GGACAGTTGAAGCCTTCCACGCCGTTACAGGCAA 3’; epitope 4: fwd 5’ AGCTTTGCCTGTGATATCCCGATCGGCGCGGGCATCTGTCCTGCA 3’, rev 5’ GGACAGATGCCCGCGCCGATCGGGATATCACAGGCAA 3’; epitope 5: fwd 5’ AGCTTTGCCTGTACCATGTATATCTGTGGCGATAGCACCGAATGTAG 3’, rev 5’ CAACCTGCTGCTGCAGTATGGCAGCTTCTGTCCTGCA 3’; epitope 6: fwd 5’ GGACAGAAGATCGCCACAGATATACATGGTACAGGCAA 3’, rev 5’ AGCTTTGCCTGTGTGCTGGGCCAGAGCAAACGCGTGGATTTCTGTCC 3’ were mixed at equimolar ratio and annealed using a thermocycler (93°C for 3 min, 80°C for 20 min, 75°C for 20 min, 70°C for 20 min, 65°C for 20 min, 40°C for 60 min). The f88-4 plasmid was digested with HindIII and PstI restriction endonucleases and ligated with the annealed double stranded oligonucleotides as described (18, 19). Each ligation product was electroporated into electrocompetent DH5α *E*.*coli*. Sequence-verified individual clones were used to infect K91kan *E*.*coli*. Phage particles were cultured in LB media containing 1 mM IPTG, tetracycline (40 μg/mL), and kanamycin (100 μg/mL), and were purified by polyethylene glycol (PEG)-NaCl precipitation method (17). Titration of single-display phage particles was carried out by infection of host bacterial cells K91kan *E. coli* for colony counting and represented as transducing units (TU/µL).

### Generation of Dual-Display Phage Particles

To produce phage particles simultaneously displaying SARS-CoV-2 S protein epitope 4 (aa 662–671) on rpVIII protein and the lung transport peptide CAKSMGDIVC on pIII protein, we fused the single-display phage constructs (described above) and the fUSE55 genome to create a chimeric vector. The f88-4 vector-derived DNA fragment containing the rpVIII gene was inserted into the fUSE55 phage vector by double digestion of both vectors with XbaI and BamHI restriction enzymes at 37 °C, 4 h. After incubation, DNA fragments were loaded onto an agarose gel (0.8%, wt/vol). Under an ultraviolet transilluminator, the 3,925 bp fUSE55 DNA fragment and the 5,402-bp f88-4 DNA fragment containing epitope 4 in-frame with the rpVIII gene were excised. Fifty nanograms (ng) of the fUSE55 DNA fragments were ligated to 68.8 ng of the f88-4 DNA fragment with T4 DNA ligase (1U) in a final volume of 20 μL, O.N. at 16 °C for 16 h. An aliquot of the ligation reaction was transformed into electrocompetent DH5α *E*.*coli* and plated onto LB agar plates containing 40 μg/mL of tetracycline. Positive clones were verified by Sanger sequencing analysis and the plasmid containing the chimeric vector was purified with a QIAprep Spin Miniprep kit (Qiagen). The chimeric vector containing the f88-4 epitope 4 (aa 662–671) and fUSE55 was then digested with SfiI at 50°C for 4 h. The digested chimera was ligated to annealed oligonucleotides encoding the CAKSMGDIVC peptide into the fUSE55 pIII gene to generate the dual-display phage vector. The titration of dual-display phage particles was carried out by infection of host bacterial cells K91kan *E*.*coli*.

### Genetic Engineering and Production of RGD4C-AAVP *S* and RGD4C-AAVP *S-null* Particles

The 3.821 kb SARS-CoV-2 S protein coding sequence (Genbank Accession number NC_045512.2) was synthesized at GeneWiz (South Plainfield, NJ) with modifications to simplify subcloning into the RGD4C-AAVP-TNF genome. The single EcoRI restriction site at 1371 bp was deleted by replacing the thymidine nucleotide at position 1374 to cytosine in the wobble codon position without changing the translated asparagine residue at position 437. The 69-nucleotide sequence of the human interferon leader sequence and the 19-nucleotide sequence of the poly A region in RGD4C-AAVP-TNF (Genbank Accession number URN.LOCAL. P-7734T13) were added to the 5’ or 3’ ends, respectively, to the modified synthetic SARS-CoV-2 S gene to produce a 3.909 kb modified SARS-CoV-2 S gene that was subcloned into the EcoRI and SalI restriction sites of pUC57/Amp^R^ at GeneWiz. The first EcoRI restriction site at 829 bp within the AgeI and KasI restriction sites in RGD4C-AAVP-TNF was deleted in two steps to mutate a thymidine to cytosine nucleotide at position 833 without altering the translated amino acid using the Q5 site-directed mutagenesis kit (New England Biolabs, Ipswich, MA) by following the manufacturer’s protocol. Positive clones were verified by EcoRI restriction mapping, confirmed with overlapping sense and antisense primers by Sanger sequencing (SeqStudio, Thermo Fisher Scientific) using the BigDye Terminator v.3.1 Cycle Sequencing and XTerminator Purification kits (Applied Biosystems, Thermo Fisher Scientific) and analyzed using SnapGene software (GSL Biotech, San Diego, CA). Positive RGD4C-AAVP-TNFΔEcoRI829/MC1061 F^-^ colonies containing a unique EcoRI site at 10,191 kb were identified by EcoRI restriction mapping of dsDNA purified from 50 mL overnight animal-free LB Broth, pH 7.4 cultures containing 100 µg/mL streptomycin (VWR, Radnor, PA) and 40 µg/mL tetracycline (Gold Biotechnology, Inc., St. Louis, MO) using the PureLink HiPure Plasmid MidiPrep kit (Invitrogen, Thermo Fisher Scientific). Sequences near the AgeI and KasI cloning sites were confirmed by Sanger sequencing in both directions as described above. The modified, synthetic SARS CoV-2 S gene was ligated into the EcoRI/SalI sites of dephosphorylated, gel-purified, RGD4C-AAVPΔEcoRI829 digested with EcoR1-HF and SalI-HF (New England Biolabs) using a 1:3 vector:insert ratio and a vector mass of 15 ng. Ligation products were transformed into animal-free, electrocompetent MC1061 F^-^ *E*.*coli* and plated onto animal-free LB agar containing 100 µg/mL streptomycin and 40 µg/mL tetracycline (MilliporeSigma). Positive clones were verified by restriction mapping and confirmed by Sanger sequencing of purified dsDNA as described above. The 84 bp transgene null sequence containing the upstream AAVP human interferon leader sequence ending with a stop codon was synthesized at IDT (San Diego, CA) as a G-block, amplified by PCR, digested with EcoRI-HF and SalI-HF, subcloned into dephosphorylated, gel purified RGD4C-AAVP-TNFΔEcoRI829 and transformed into electrocompetent MC1061 F^-^ to produce the RGD4C-AAVP-transgene null genome. Transformed colonies were screened by colony PCR and restriction mapping. The transgene null insert was verified in putative positive clones by Sanger sequencing in both directions as described above. A single transformed RGD4C-AAVP SARS CoV-2 *S*/MC1061 F^-^ or RGD4C-AAVP SARS-CoV-2 *S-null*/MC1061 F^-^ colony was used to inoculate 10 mL of animal-free LB Broth, pH 7.4 containing 100 µg/mL streptomycin and 40 µg/mL tetracycline and grown to mid-log phase at 37 °C and 250 rpm in the dark. One milliliter of each mid-log phase pre-culture was used to inoculate 750 mL of animal-free LB Broth, pH 7.4 containing 100 µg/mL streptomycin and 40 µg/mL tetracycline in a sterile 2 L shaker baffle flasks for phage amplification at 30 °C, 250 rpm for 20 h in the dark. Phage were precipitated twice in sterile 16.7% PEG 8000/3.3 M NaCl (15% v/v), and the final phage pellet was resuspended in 1 mL sterile phosphate-buffered saline, pH 7.4, centrifuged to remove residual bacterial debris and filtered sterilized through a 0.2 µm syringe filter. Infective phage titers [(TU)/µL] were determined by infecting K91kan *E*.*coli* bacteria grown in animal-free terrific broth with 1E10^7^, 1E10^8^ or 1E10^9^ diluted phage, plating the infected bacteria onto animal-free LB agar plates containing 100 µg/mL kanamycin sulfate and 40 µg/mL tetracycline and counting bacterial colonies the following day.

### Immunization Studies in Mice

Swiss Webster or BALB/C mice were randomized in groups of 3 to 12 animals. Group size was calculated based on statistical considerations to yield sufficient statistical significance. The animals were inoculated with 10^9^ TU phage or AAVP constructs intraperitoneally, intravenously, intratracheally or subcutaneously. For subcutaneous injections, 10^9^ TU phage or AAVP particles were administered with 100 μL on the front and hind limbs, and behind the neck (∼ 20 μL per site). For intratracheal vaccination, 10^9^ TU of single-, dual-display phage particles or negative control insertless phage particles were administered in two serial doses in 50 μL of PBS with a MicroSprayer® Aerosolizer coupled to a high-pressure syringe (Penn-Century) and a small animal laryngoscope (Penn-Century). The devices were used to administer air-free liquid aerosol directly into the trachea of animals deeply anesthetized with 1% isoflurane (22). For the tail vein blood collections, mice were locally anesthetized with a topic solution. On day 0, blood samples were collected for the baseline, followed by consecutive blood collection every 1-2 weeks post-immunization. Endotoxin removal was performed for each phage or AAVP preparation prior to administration of each dose, regardless of the route of administration. Purified phage or AAVP containing endotoxin was mixed with 10% Triton X-114 in endotoxin-free water, incubated on ice for 10 min, warmed to 37°C degrees for 10 min followed by separation of the Triton X-114 phase by centrifugation at 14,000 rpm for 1 min. The upper aqueous phase containing phage was withdrawn into a sterile microcentrifuge tube. This process was repeated from 3-5 times followed by PEG-NaCl precipitation. Resuspended phage was sterile filtered through 0.2 μm (Pall Corporation) syringe filters. The levels of endotoxin were measured using the Limulus Amebocyte Lysate (LAL) Kinetic-QCL kit (Lonza). Phage or AAVP preparations containing endotoxin levels < 0.05 EU/mL were used in this study.

### Serological Analysis

ELISA assays were performed in 96-microwell plates coated with 150 ng/well of SARS-CoV-2 Spike (aa 16-1213) His-tagged recombinant protein (ThermoFisher) and 10^10^ phage or AAVP particles/50 μL of PBS O.N. at 4°C (Nunc MaxiSorp flat bottom, ThermoFisher Scientific). Coated plates were blocked with PBS containing 5% low-fat milk and 1% BSA for 1 h at 37°C. Two-fold serial dilutions (starting at 1:32) or 1:50 fixed dilution of sera in blocking buffer were added to separate the wells and incubated for 1-2 h at 37°C. Following three washes with PBS and PBS containing 0.1% of Tween 20, bound antibodies were detected with an anti-mouse IgG HRP-conjugated (Jackson ImmunoResearch) at optical density (OD) at 450 nm. Commercially available polyclonal IgG anti-Spike protein antibody (Thermo Fisher, MA5-35949) or anti-fd bacteriophage antibody (Sigma Aldrich) served as positive controls.

### RNA Isolation and Quantitative Real-Time PCR

Total RNA from mice tissues were obtained with the RNeasy Mini Kit (Qiagen). First-strand cDNA synthesis was carried out with the ImProm-II Reverse Transcription System (Promega). Quantitative real-time PCR analysis was performed in a QuantStudio 5 Real-Time PCR System (Applied Biosystems). Primers and TaqMan probes were as follows: fwd 5’ GCTTTTCAGCTCTGCATCGTT 3’ and rev 5’ GACTAGTGGCAATAAAACAAGAAAAACA 3, customized AAVP *S* 6FAM 5’ TGGGTTCTCTTGGCATGT 3’ NFQ, Mm04277571_s1 for 18S, and Mm99999915_g1 for Gapdh. The gene expression ratio was normalized to that of 18S.

### Statistical Analysis

Differences between groups were tested for statistical significance with Student’s *t*-test or analysis of variance (one-way or two-way ANOVA) using GraphPad Prism 8. Statistical significance was set at *P* < 0.05.

## Acknowledgments

The authors wish to thank Dr. Helen Pickersgill (Life Science Editors) for professional editorial services and Dr. Isan Chen and Mr. Jason Rifkin for the critical reading of the manuscript. This work was supported in part by core services from Rutgers Cancer Institute of New Jersey Cancer Center Support Grant (P30CA072720); by the RCSB Protein Data Bank, which is jointly funded by the National Science Foundation (DBI-1832184), the US Department of Energy (DE-SC0019749), and the National Cancer Institute, National Institute of Allergy and Infectious Diseases, and National Institute of General Medical Sciences of the National Institutes of Health (R01GM133198); by the National Science Foundation (NSF) grants (CHE-1614101 and PHY-1522550 to J.N.O. and MCB-1915843 to P.C.W), and by research awards from the Gillson-Longenbaugh Foundation (to R.P. and W.A.) and Welch Foundation (C-1792) (to J.N.O.). The work at the Center for Theoretical Biological Physics was also supported by the NSF (Grant PHY-2019745). J.N.O. is a Cancer Prevention Research in Texas Scholar in Cancer Research. The authors also wish to acknowledge generous support from the Northeastern University Discovery cluster and Northeastern University Research Computing staff.

## Author contribution

D.I.S., F.H.F.T., C.M., V.J.Y., F.I.S., E.D.R.,V.G.C., D.D., P.O’B., N.H., T.L.S., R.L.S., P.C.W., J.N.O., W.A., and R.P. designed the research; D.I.S., F.H.F.T., C.M., V.J.Y., F.I.S., E.D.R., V.G.C., D.D., P.O’B., N.H., T.L.S., N.B., M.L.G. and P.C.W. performed the research; E.C.L., S.K.L., S.K.B., J.N.O. contributed new reagents/analytic tools; D.I.S., F.H.F.T., C.M., V.J.Y., F.I.S., E.D.R., V.G.C., D.D., P.O’B., N.H., T.L.S., N.B., R.L.S., M.L.G., P.C.W., J.N.O, W.A. and R.P analyzed the data; D.I.S., F.H.F.T., C.M., V.J.Y., F.I.S., V.G.C., E.D.R., P.C.W, J.N.O., W.A. and R.P. wrote the initial draft of the manuscript, to which all of the authors contributed with edits. J.N.O., W.A. and R.P. supervised the overall project.

## Supplemental Figures

**Fig S1.**
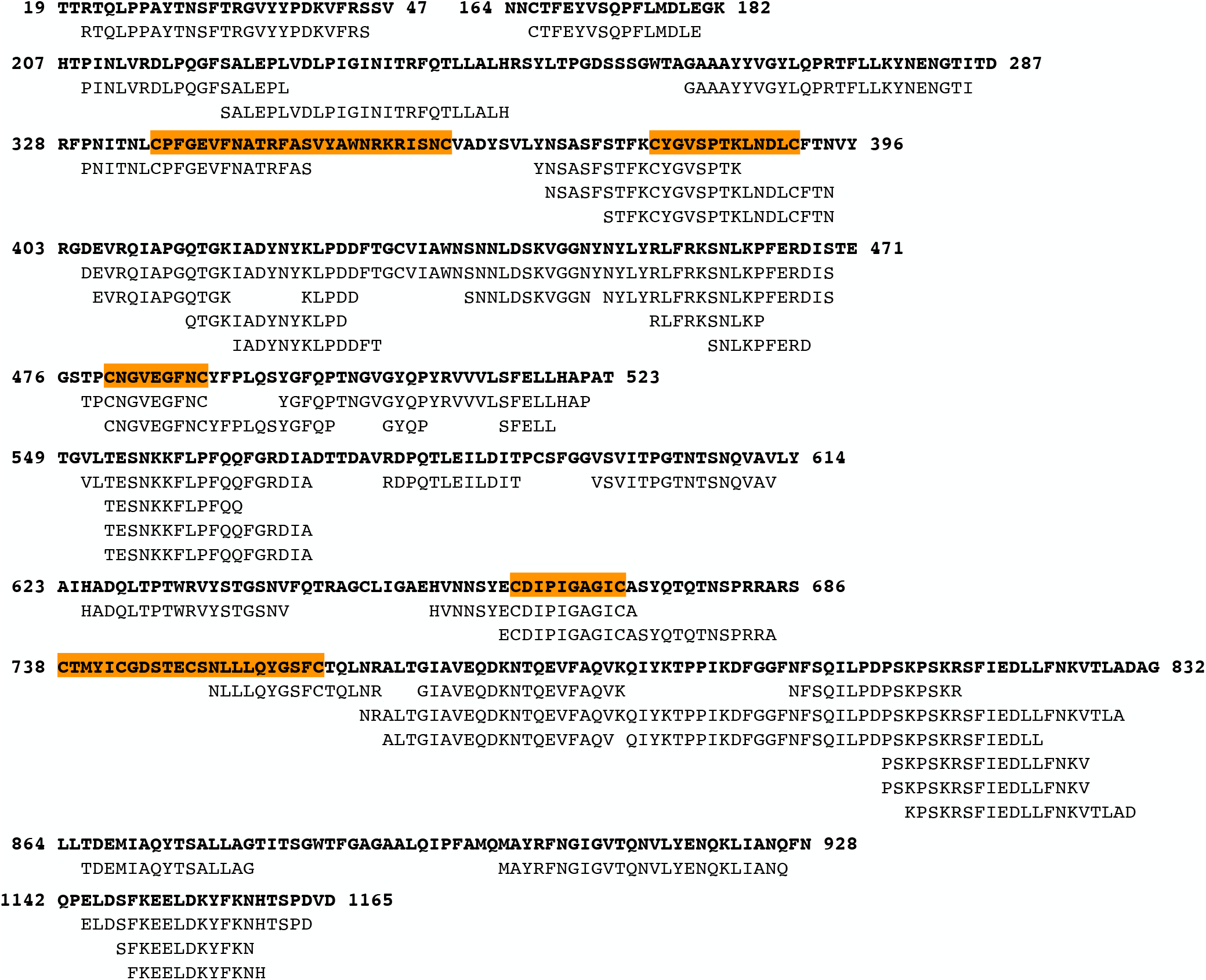
S protein epitope mapping from peer-reviewed publications. Primary sequence representation of immunogenic regions of SARS-CoV-2 S protein spanning S1 and S2 subunits identified by B-cells, T-cells, and antibody screenings on patients with COVID-19 (also listed in Table S1). The six structurally-selected epitopes displayed on the rpVIII protein are highlighted (orange).

**Fig S2.**
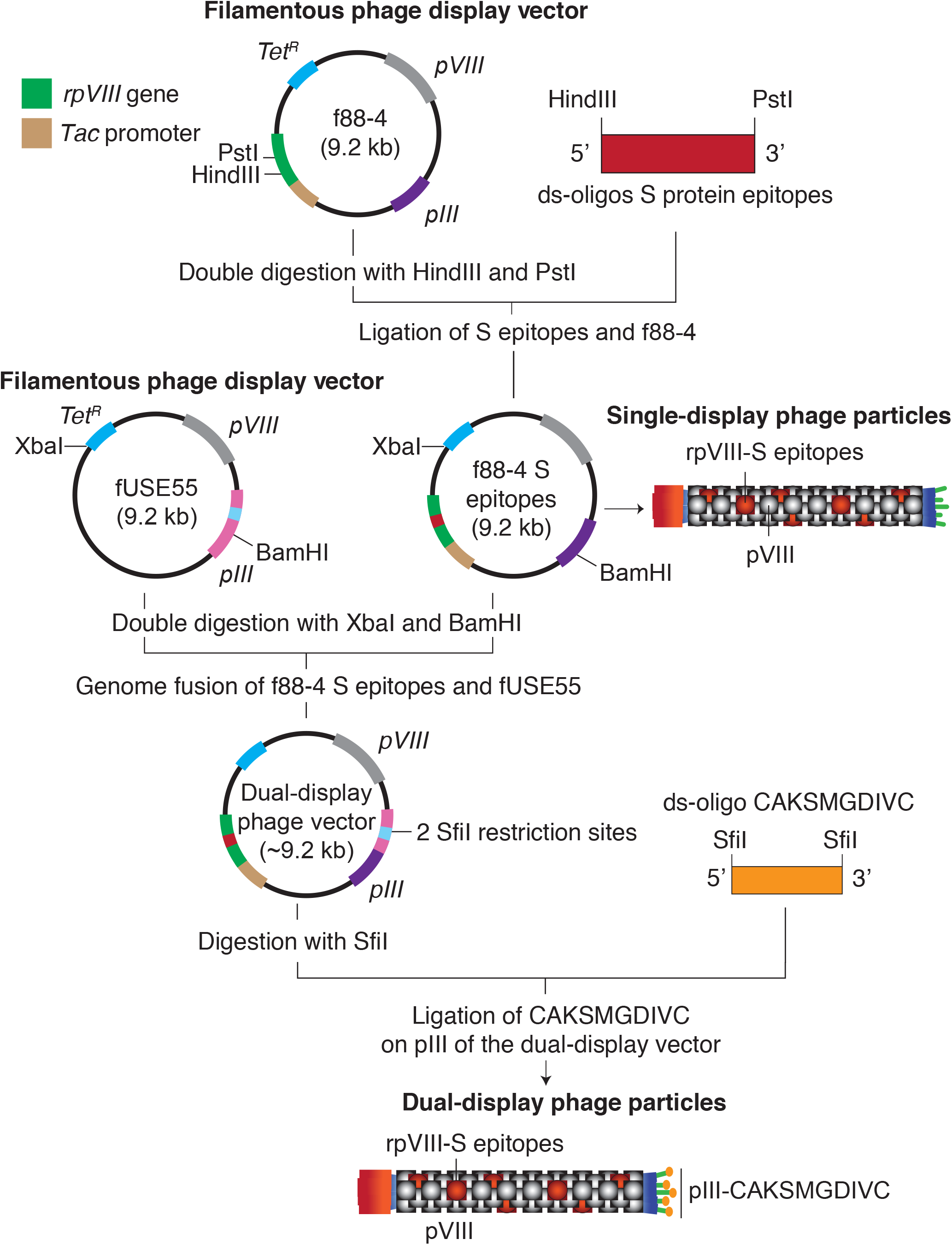
The single- and dual-display phage particles cloning strategy. To generate the single-display phage particles, the f88-4 phage vector was used. This vector contains two genes encoding for the major capsid protein pVIII: the wild-type (pVIII, depicted in grey) and the recombinant (rpVIII, depicted in green). rpVIII contains a foreign DNA insert between the HindIII and PstI cloning site, which allows the cloning of annealed oligonucleotides encoding the S protein epitopes in-frame with the rpVIII gene. For the dual-display phage particles, the f88-4 vector containing the epitope 4 (CDIPIGAGIC) and the fUSE55 phage vector are digested with BamHI and XbaI restriction enzymes. The digestion products are then purified and fused according to a standard ligation protocol. The result is a chimeric vector (f88-4/fUSE55). Next, oligonucleotide encoding for the CAKSMGDIVC targeting motif was cloned within the SfiI restriction site of the pIII gene (pIII, depicted in light blue), generating the dual-display phage vector, which contains epitope 4 on the rpVIII protein and the CAKSMGDIVC motif on the pIII protein. Both, single-or dual-display phage particle vectors were used to transform electrocompetent DH5α *E. coli*. Phage particles were produced in k91Kan *E. coli*. Tet^R^, tetracycline resistance gene.

---

**Table S1:**
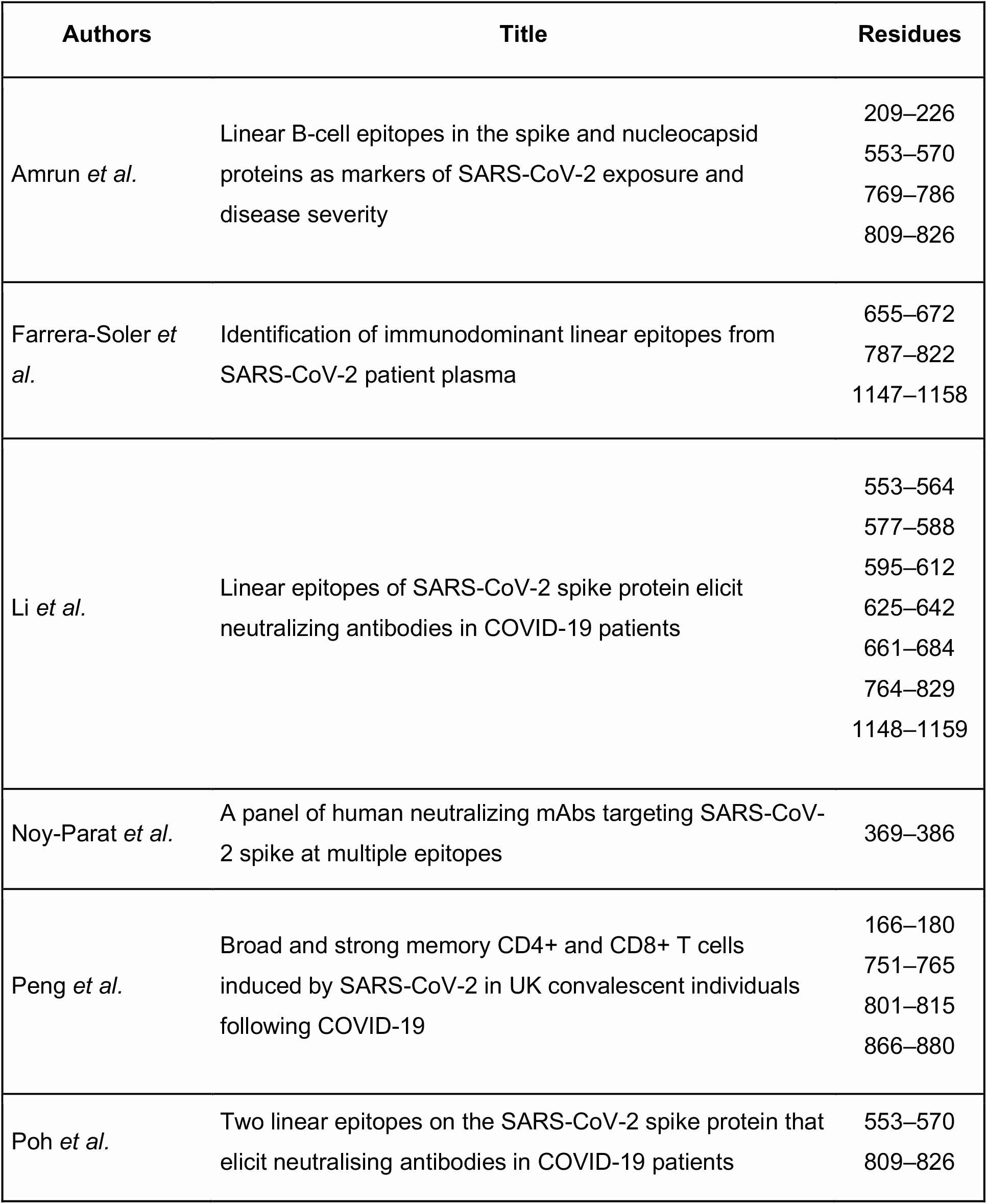
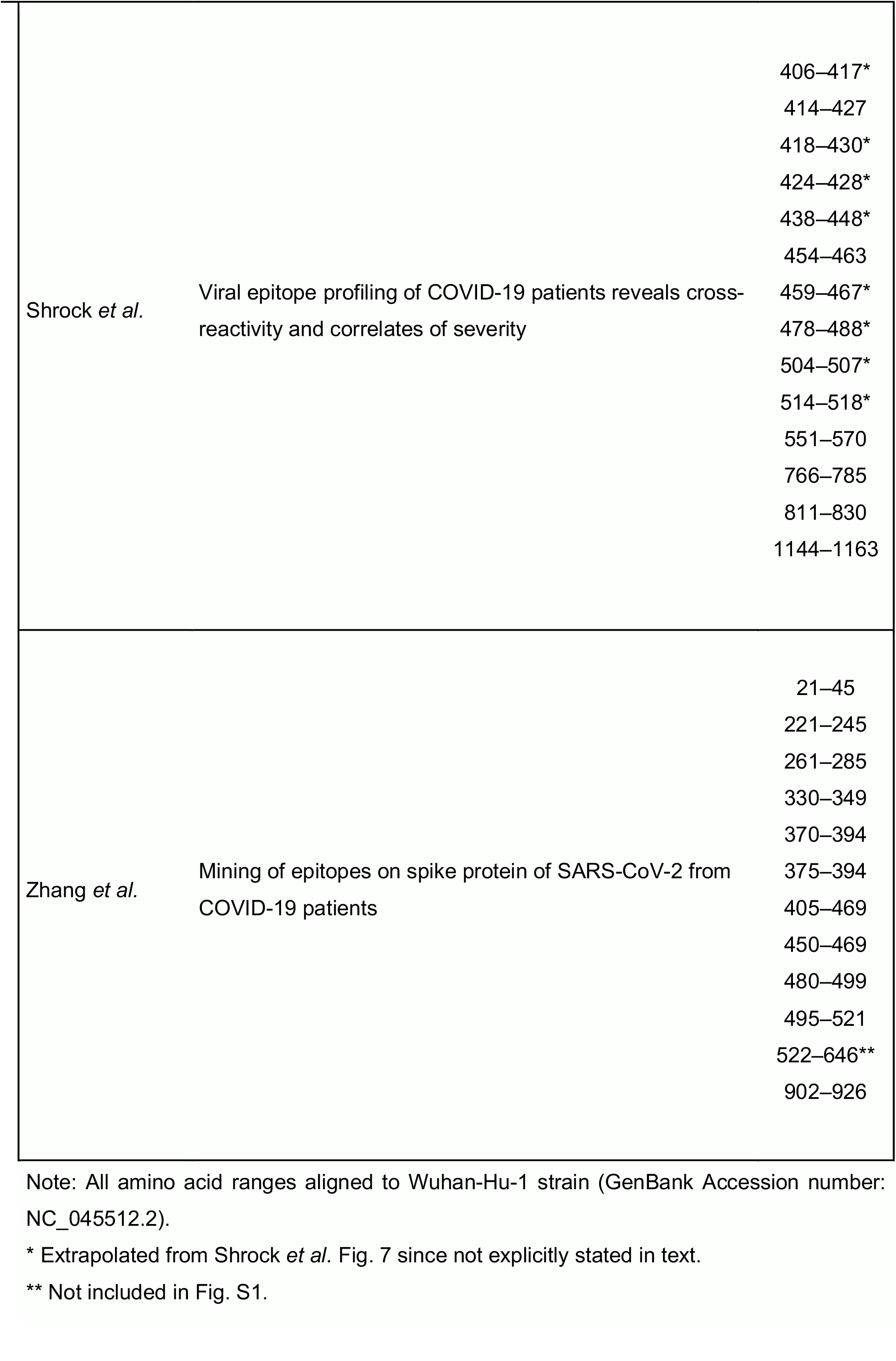
Cross-reference analysis of epitope mapping of immunogenic regions of SARS-CoV-2 S protein.

## References

1. F. Wu et al., A new coronavirus associated with human respiratory disease in China. Nature 579, 265–269 (2020).

2. P. Zhou et al., A pneumonia outbreak associated with a new coronavirus of probable bat origin. Nature 579, 270–273 (2020).

3. Y. Huang, C. Yang, X. F. Xu, W. Xu, S. W. Liu, Structural and functional properties of SARS-CoV-2 spike protein: potential antivirus drug development for COVID-19. Acta Pharmacol Sin 41, 1141–1149 (2020).

4. A. C. Walls et al., Structure, function, and antigenicity of the SARS-CoV-2 Spike glycoprotein. Cell 181, 281–292 e286 (2020).

5. A. Baum et al., REGN-COV2 antibodies prevent and treat SARS-CoV-2 infection in rhesus macaques and hamsters. Science 370, 1110–1115 (2020).

6. B. Korber et al., Tracking changes in SARS-CoV-2 Spike: evidence that D614G increases infectivity of the COVID-19 virus. Cell 182, 812–827 e819 (2020).

7. Q. Li et al., The impact of mutations in SARS-CoV-2 Spike on viral infectivity and antigenicity. Cell 182, 1284–1294 e1289 (2020).

8. S. T. Abedon, S. J. Kuhl, B. G. Blasdel, E. M. Kutter, Phage treatment of human infections. Bacteriophage 1, 66–85 (2011).

9. E. M. Barbu, K. C. Cady, B. Hubby, Phage therapy in the era of synthetic biology. Cold Spring Harb Perspect Biol 8 (2016).

10. A. Gonzalez-Mora, J. Hernandez-Perez, H. M. N. Iqbal, M. Rito-Palomares, J. Benavides, Bacteriophage-based vaccines: A potent approach for antigen delivery. Vaccines (Basel) 8 (2020).

11. K. A. Henry, M. Arbabi-Ghahroudi, J. K. Scott, Beyond phage display: non-traditional applications of the filamentous bacteriophage as a vaccine carrier, therapeutic biologic, and bioconjugation scaffold. Front Microbiol 6, 755 (2015).

12. M. Trepel, W. Arap, R. Pasqualini, Modulation of the immune response by systemic targeting of antigens to lymph nodes. Cancer Res 61, 8110–8112 (2001).

13. W. Arap et al., Steps toward mapping the human vasculature by phage display. Nat Med 8, 121–127 (2002).

14. R. D. Pentz et al., Ethics guidelines for research with the recently dead. Nat Med 11, 1145–1149 (2005).

15. F. I. Staquicini et al., Vascular ligand-receptor mapping by direct combinatorial selection in cancer patients. Proc Natl Acad Sci U S A 108, 18637–18642 (2011).

16. R. Pasqualini, E. Ruoslahti, Organ targeting in vivo using phage display peptide libraries. Nature 380, 364–366 (1996).

17. R. Pasqualini, E. Koivunen, E. Ruoslahti, Alpha v integrins as receptors for tumor targeting by circulating ligands. Nat Biotechnol 15, 542–546 (1997).

18. R. Rangel et al., Combinatorial targeting and discovery of ligand-receptors in organelles of mammalian cells. Nat Commun 3, 788 (2012).

19. R. Rangel et al., Targeting mammalian organelles with internalizing phage (iPhage) libraries. Nat Protoc 8, 1916–1939 (2013).

20. A. Hajitou et al., A hybrid vector for ligand-directed tumor targeting and molecular imaging. Cell 125, 385–398 (2006).

21. K. Suwan et al., Next-generation of targeted AAVP vectors for systemic transgene delivery against cancer. Proc Natl Acad Sci U S A 116, 18571–18577 (2019).

22. D. I. Staquicini et al., Targeted phage display-based pulmonary vaccination in mice and non-human primates. Med 3 (2020).

23. M. C. Paoloni et al., Launching a novel preclinical infrastructure: comparative oncology trials consortium directed therapeutic targeting of TNFalpha to cancer vasculature. PLoS One 4, e4972 (2009).

24. F. I. Staquicini et al., Systemic combinatorial peptide selection yields a non-canonical iron-mimicry mechanism for targeting tumors in a mouse model of human glioblastoma. J Clin Invest 121, 161–173 (2011).

25. T. L. Smith et al., AAVP displaying octreotide for ligand-directed therapeutic transgene delivery in neuroendocrine tumors of the pancreas. Proc Natl Acad Sci U S A 113, 2466–2471 (2016).

26. A. S. Dobroff et al., Towards a transcriptome-based theranostic platform for unfavorable breast cancer phenotypes. Proc Natl Acad Sci U S A 113, 12780–12785 (2016).

27. F. Ferrara et al., Targeted molecular-genetic imaging and ligand-directed therapy in aggressive variant prostate cancer. Proc Natl Acad Sci U S A 113, 12786–12791 (2016).

28. F. I. Staquicini et al., Targeted AAVP-based therapy in a mouse model of human glioblastoma: a comparison of cytotoxic versus suicide gene delivery strategies. Cancer Gene Ther 27, 301–310 (2020).

29. K. Dijkman et al., Prevention of tuberculosis infection and disease by local BCG in repeatedly exposed rhesus macaques. Nat Med 25, 255–262 (2019).

30. N. B. Mercado et al., Single-shot Ad26 vaccine protects against SARS-CoV-2 in rhesus macaques. Nature 586, 583–588 (2020).

31. M. Meyer et al., Aerosolized Ebola vaccine protects primates and elicits lung-resident T cell responses. J Clin Invest 125, 3241–3255 (2015).

32. A. O. Hassan et al., A single-dose intranasal ChAd vaccine protects upper and lower respiratory tracts against SARS-CoV-2. Cell 183, 169–184 e113 (2020).

33. E. Koivunen, B. Wang, E. Ruoslahti, Phage libraries displaying cyclic peptides with different ring sizes: ligand specificities of the RGD-directed integrins. Biotechnology (N Y) 13, 265–270 (1995).

34. M. G. Overstreet et al., Inflammation-induced interstitial migration of effector CD4(+) T cells is dependent on integrin alphaV. Nat Immunol 14, 949–958 (2013).

35. N. Okada et al., Dendritic cells transduced with gp100 gene by RGD fiber-mutant adenovirus vectors are highly efficacious in generating anti-B16BL6 melanoma immunity in mice. Gene Ther 10, 1891–1902 (2003).

36. A. Yano et al., RGD motif enhances immunogenicity and adjuvanicity of peptide antigens following intranasal immunization. Vaccine 22, 237–243 (2003).

37. W. P. Poh et al., Characterization of cytotoxic T-lymphocyte epitopes and immune responses to SARS coronavirus spike DNA vaccine expressing the RGD-integrin-binding motif. J Med Virol 81, 1131–1139 (2009).

38. S. N. Amrun et al., Linear B-cell epitopes in the spike and nucleocapsid proteins as markers of SARS-CoV-2 exposure and disease severity. EBioMedicine 58, 102911 (2020).

39. C. M. Poh et al., Two linear epitopes on the SARS-CoV-2 spike protein that elicit neutralising antibodies in COVID-19 patients. Nat Commun 11, 2806 (2020).

40. L. Farrera-Soler et al., Identification of immunodominant linear epitopes from SARS-CoV-2 patient plasma. PLoS One 15, e0238089 (2020).

41. Y. Li et al., Linear epitopes of SARS-CoV-2 spike protein elicit neutralizing antibodies in COVID-19 patients. Cell Mol Immunol 17, 1095–1097 (2020).

42. T. Noy-Porat et al., A panel of human neutralizing mAbs targeting SARS-CoV-2 spike at multiple epitopes. Nat Commun 11, 4303 (2020).

43. Y. Peng et al., Broad and strong memory CD4(+) and CD8(+) T cells induced by SARS-CoV-2 in UK convalescent individuals following COVID-19. Nat Immunol 21, 1336–1345 (2020).

44. E. Shrock et al., Viral epitope profiling of COVID-19 patients reveals cross-reactivity and correlates of severity. Science 370 (2020).

45. B. Z. Zhang et al., Mining of epitopes on spike protein of SARS-CoV-2 from COVID-19 patients. Cell Res 30, 702–704 (2020).

46. S. Kumar, V. K. Maurya, A. K. Prasad, M. L. B. Bhatt, S. K. Saxena, Structural, glycosylation and antigenic variation between 2019 novel coronavirus (2019-nCoV) and SARS coronavirus (SARS-CoV). Virusdisease 31, 13–21 (2020).

47. D. Wrapp et al., Cryo-EM structure of the 2019-nCoV spike in the prefusion conformation. Science 367, 1260–1263 (2020).

48. R. L. Sheets et al., Biodistribution and toxicological safety of adenovirus type 5 and type 35 vectored vaccines against human immunodeficiency virus-1 (HIV-1), Ebola, or Marburg are similar despite differing adenovirus serotype vector, manufacturer’s construct, or gene inserts. J Immunotoxicol 5, 315–335 (2008).

49. R. L. Sheets et al., Toxicological safety evaluation of DNA plasmid vaccines against HIV-1, Ebola, Severe Acute Respiratory Syndrome, or West Nile virus is similar despite differing plasmid backbones or gene-inserts. Toxicol Sci 91, 620–630 (2006).

50. S. F. Ahmed, A. A. Quadeer, M. R. McKay, Preliminary identification of potential vaccine targets for the COVID-19 coronavirus (SARS-CoV-2) based on SARS-CoV immunological studies. Viruses 12 (2020).

51. J. L. Slon-Campos et al., A protective Zika virus E-dimer-based subunit vaccine engineered to abrogate antibody-dependent enhancement of dengue infection. Nat Immunol 20, 1291–1298 (2019).

52. N. van Doremalen et al., ChAdOx1 nCoV-19 vaccine prevents SARS-CoV-2 pneumonia in rhesus macaques. Nature 586, 578–582 (2020).

53. V. J. Munster et al., Respiratory disease and virus shedding in rhesus macaques inoculated with SARS-CoV-2. bioRxiv 10.1101/2020.03.21.001628 (2020).

54. L. Feng et al., An adenovirus-vectored COVID-19 vaccine confers protection from SARS-COV-2 challenge in rhesus macaques. Nat Commun 11, 4207 (2020).

55. L. Makowski, W. Olson-Sidford, W. W. J, Biological and clinical consequences of integrin binding via a rogue RGD motif in the SARS CoV-2 spike protein. Viruses 13 (2021).

56. D. R. Christianson et al., Ligand-directed targeting of lymphatic vessels uncovers mechanistic insights in melanoma metastasis. Proc Natl Acad Sci U S A 112, 2521–2526 (2015).

57. E. F. Pettersen et al., UCSF Chimera--a visualization system for exploratory research and analysis. J Comput Chem 25, 1605–1612 (2004).

58. I. M. Ibrahim, D. H. Abdelmalek, M. E. Elshahat, A. A. Elfiky, COVID-19 spike-host cell receptor GRP78 binding site prediction. J Infect 80, 554–562 (2020).

59. X. Xiong et al., A thermostable, closed SARS-CoV-2 spike protein trimer. Nat Struct Mol Biol 27, 934–941 (2020).

60. A. G. Wrobel et al., SARS-CoV-2 and bat RaTG13 spike glycoprotein structures inform on virus evolution and furin-cleavage effects. Nat Struct Mol Biol 27, 763–767 (2020).

61. B. Hess, C. Kutzner, D. van der Spoel, E. Lindahl, GROMACS 4: Algorithms for highly efficient, load-balanced, and scalable molecular simulation. J Chem Theory Comput 4, 435–447 (2008).

62. E. Lindahl, B. Hess, D. van der Spoel, GROMACS 3.0: a package for molecular simulation and trajectory analysis. Molecular modeling annual 7, 306–317 (2001).

63. W. G. Hoover, Canonical dynamics: Equilibrium phase-space distributions. Physical Review A 31, 1695–1697 (1985).

64. W. L. Jorgensen, J. Chandrasekhar, J. D. Madura, R. W. Impey, M. L. Klein, Comparison of simple potential functions for simulating liquid water. Journal of Chemical Physics 79, 926 (1983).

65. K. Lindorff-Larsen et al., Improved side-chain torsion potentials for the Amber ff99SB protein force field. Proteins 78, 1950–1958 (2010).

66. S. Nosé, A unified formulation of the constant temperature molecular dynamics methods. The Journal of Chemical Physics 81, 511–519 (1984).

67. M. Parrinello, A. Rahman, Polymorphic transitions in single crystals: A new molecular dynamics method. Journal of Applied Physics 52, 7182 (1981).

